# Gorse (*Ulex europeaus*) wastes with 5,6-dimethyl benzimidazole supplementation can support growth of vitamin B_12_ producing commensal gut microbes

**DOI:** 10.1101/2023.08.02.551616

**Authors:** Ajay Iyer, Eva C. Soto-Martín, Gary A. Cameron, Petra Louis, Sylvia H. Duncan, Charles S. Bestwick, Wendy R. Russell

## Abstract

Many commensal gut microbes are recognized for their potential to synthesize vitamin B_12_, offering a promising avenue to address deficiencies through probiotic supplementation. While bioinformatics tools aid in predicting B_12_ biosynthetic potential, empirical validation remains crucial to confirm production, identify cobalamin vitamers, and establish biosynthetic yields.

This study investigates vitamin B_12_ production in three human colonic bacterial species: *Anaerobutyricum hallii* DSM 3353, *Roseburia faecis* DSM 16840, and *Anaerostipes caccae* DSM 14662, along with *Propionibacterium freudenreichii* DSM 4902 as a positive control. These strains were selected for their potential use as probiotics, based on speculated B_12_ production from prior bioinformatic analyses. Cultures were grown in M2GSC, chemically defined media (CDM), and Gorse extract medium (GEM). The composition of GEM was similar to CDM, expect that the carbon and nitrogen source was replaced with protein-depleted liquid wastes obtained after subjecting Gorse to a leaf protein extraction process. B_12_ yields were quantified using liquid chromatography with tandem mass spectrophotometry.

The results suggest that the three butyrate-producing strains could indeed produce B_12_, although the yields were notably low, and with no B_12_ being detected in the extracellular media. Next, GEM outperformed the conventional M2GSC growth medium in enhancing B_12_ production. The positive control, *P. freudenreichii* DSM 4902 produced B_12_ at concentrations ranging from 7 ng·mL^−1^ to 12 ng·mL^−1^. Univariate-scaled Principal Component Analysis of data from previous publications investigating B_12_ production in *P. freudenreichii* revealed that B_12_ yields diminished when the carbon source concentration was *≤*30 g·L^−1^.

In conclusion, the protein-depleted wastes from the leaf protein extraction process from Gorse can be valorised as a viable substrate for culturing B_12_-producing colonic gut microbes. Furthermore, this is the first report attesting to the ability of *A. hallii, R. faecis*, and *A. caccae* to produce B_12_. However, these microbes seem unsuitable for industrial applications owing to low B_12_ yields.

## Introduction

### The biological role and molecular structure of vitamin B_12_

Vitamin B_12_ is essential for various enzymatic reactions, including isomerases, methyltransferases, and dehalogenases (Banerjee & Ragsdale, 2003), with B_12_-dependent dehalogenases being exclusive to bacteria (Payne et al., 2015). It is a critical cofactor in the type 1 reaction, catalysing the isomerization of L-methylmalonyl-CoA to succinyl-CoA, that subsequently enters the Krebs cycle (Takahashi-Iñiguez et al., 2012). Furthermore, B_12_ is crucial for the type 2 reaction of methyl transfer from 5-methyltetrahydrofolate to homocysteine, to produce tetrahydrofolate and methionine (Froese et al., 2019). Hindrance in this type 2 reaction owing to B_12_ deficiency results in the malformation of red blood corpuscles, and manifests as pernicious anaemia (World Health Organization & Food and Agriculture Organization of the United Nations, 2004).

The bioactivity of vitamin B_12_ hinges on three critical ligands, namely, the central cobalt ion located in the upper corrin ring, the methyl side-group, and 5,6-dimethylbenzimidazole (DMB) (see Figure 1). The methyl side-group and DMB are attached to the cobalt ion with a coordinate bond. The cobalt ion switches from the oxidised intermediate (III) to the reduced (I) state during the transfer of the methyl-group to homocysteine to form methionine. The oxidised cobalamin is subsequently re-alkylated with n-methyl-THF. The exact mechanism has been described previously by Matthews et al. (2008).

**Figure 1.**
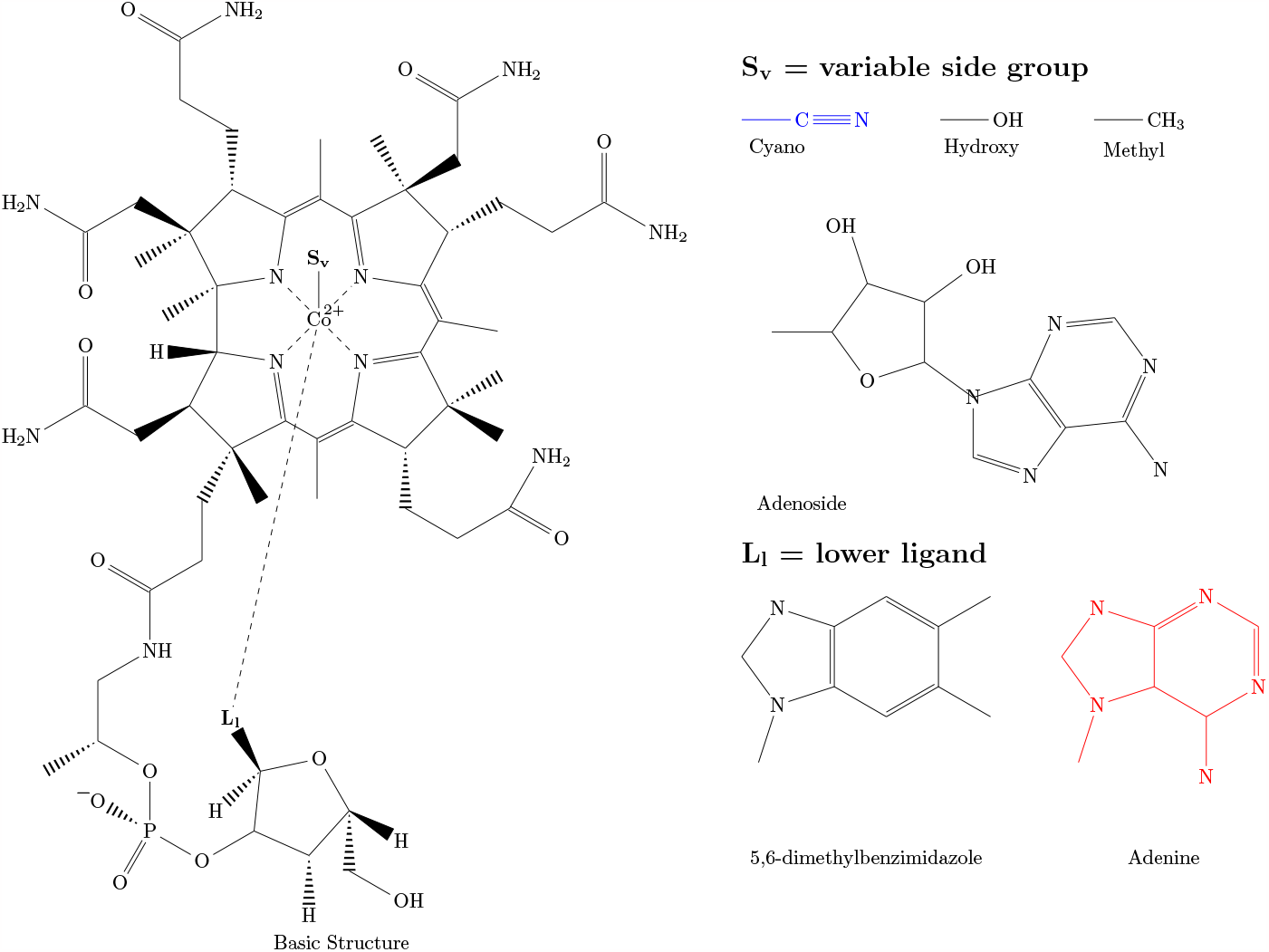
The variable ligands of B_12_ vitamers. Variable side-groups form coordinate bonds with the central cobalt ion. The cyano-group is marked in blue as it is a semi-synthetic derivate of B_12_. The ligands at location L_l_ determine if the cobalamin vitamer is biologically active in humans with 5,6-dimethybenzimidazole (DMB) or inactive with adenine (marked in red). B_12_ vitamers can be found with combinations of S_v_ and L_l_ ligands.

Dietary vitamin B_12_ however may have either the methyl, adenosyl, hydroxyl or cyano group associated with the cobalt atom, with the cyano vitamer being a semi-synthetic stable derivative of B_12_ (Herbert, 1988). Once absorbed in the body, the molecule is converted to the hydroxyl-form, that is then re-alkylated to the methyl form; ultimately entering the methionine synthesis pathway as described previously (Froese et al., 2019). There is variation in the second ligand moiety, where either the biologically active DMB, or its structural homologue, adenyl-ribofuranose phosphate, may be present. Only the DMB-based vitamer is capable of participating in the catalytic reaction facilitated by methionine synthase.

### Harnessing microbes for industrial B_12_ production

The biosynthesis of B_12_ requires several enzymes, that are only found in the bacteria and archaea domains of life (Raux et al., 2000; Stamford et al., 1997). The complete set of genes required for *de novo* production of cobalamins; particularly B_12_, has been observed in numerous bacteria such as *Pseudomonas dentrificans, Bacillus megaterium, Rhodobacter capusulatus, Bacillus megaterium* and *Thermosipho melanesiensis*, to name a few (Fang et al., 2017). Many other bacteria such as *Escherichia coli, Kosmotoga olearia* and *Fervidobacterium nodosum* can only use salvage pathways to synthesise cobalamin from precursors. The population of these microbes capable of cobalamin production thought salvage pathways play an important role in the micro-ecology of the niche they inhabit firstly through commensalistic or even symbiotic association with primary B_12_ producers (Raux et al., 2000), and secondly, by facilitating tertiary associations with other microbes that are heterotrophic for the precursor. In industrial setups however, candidate selection of B_12_ producers is largely influenced by the genetic stability of the strain, bioconversion efficiency, B_12_ yields, extracellular production capability, and rapid turnover.

Many microbial species; either naturally, or through genetic engineering, are capable of over-synthesising vitamin B_12_. The most commonly used species in industry are *Propionibacterium freudenreichii, Propionibacterium shermanii* and *Pseudomonas denitrificans*. Growth conditions are generally aerobic as *de novo* biosynthesis of vitamin B_12_ is an energetically demanding process, particularly the synthesis of DMB (Raux et al., 2000).

Consequently, there is great variability in the growth substrates used, depending on the metabolic capability of the microbe employed. For example, many media formulations use additional supple-mentation of cobalt, DMB, and in rare cases, haeme (Chen & Wolin, 1981). This helps cope with the increased demand for micronutrients, and alleviates the metabolic burden of *de novo* DMB or corrin synthesis. Interestingly, this has lead to investigations on waste stream valorisation with cobalt and DMB supplementation for the growth of B_12_ producing bacteria. For example, *P. freudenreichii* has been shown to grow on apple pomace and potato wastes for large scale production of propionic acid, with vitamin B_12_ obtained as a secondary product in significant quantities (Piwowarek et al., 2022; Piwowarek et al., 2023). Overall, such endeavours help generate high-value products and circularise waste streams otherwise fated for landfill or waste-water treatment plants.

### B_12_ from Gorse wastes using gut microbes

The investigation reported herein attempts to valorise the liquid wastes generated from a previously described leaf protein extraction process, that was obtained from the biomass of Gorse (*Ulex europaeus*), which has become invasive in many parts of the world (Iyer et al., 2022). The process generated a protein depleted fraction (PDF) that contained residual proteins, plant bioactives, and carbohydrates. In an effort to valorise this waste stream, three butyrate-producing microbial candidates, namely *Anaerobutyricum hallii* DSM 3353, *Roseburia faecis* DSM 16840, and *Anaerostipes caccae* DSM 14662, were investigated for their ability to grow on media comprising of PDF as the sole carbon and nitrogen source. These microbial candidates were chosen due to their potential to produce vitamin B_12_ (Soto-Martín et al., 2020) and their purported beneficial impact in the human gut (Arruda et al., 2022; Montalban-Arques et al., 2021). *Propionibacterium freudenreichii* DSM 4902, was used as a positive control for vitamin B_12_ production.

## Materials and methods

All chemicals were purchased from Merck, (Darmstadt, Germany) unless stated otherwise. All water used was from the in-house Milli-Q^®^purification system. All experiments were performed in triplicate.

### Media

Media preparation was carried out in an anaerobic cabinet (Don Whitley Scientific, West Yorkshire, UK) set for 10% H_2_, 10% CO_2_, and 80% N_2_, with Milli-Q water which was first flushed with N_2_ and autoclaved. The water was then allowed to equilibrate in the cabinet prior to preparation for 48 h.

### Media preparation

#### M2GSC

The recipe for M2GSC is provided in **S1 Text**, and was prepared as described previously by Cummings and Macfarlane (1991). The medium was allowed to equilibrate in the anaerobic cabinet for 48 h before use.

#### Chemically defined medium (CDM)

CDM was prepared as described previously by Soto-Martín et al. (2020), except with the inclusion of 5,6-dimethylbenzimidazole (DMB, 20 mg·L^−1^). The medium was filter-sterilised using 0.2 μm filter (Millipore, Merck, US) and allowed to equilibrate in the anaerobic cabinet. The composition of the CDM is provided in **S2 Text**.

#### Gorse extract medium (GEM)

GEM was prepared by replacing the carbon and nitrogen source of CDM with the gorse protein-depleted fraction (PDF). The composition of PDF is provided in a previous publication by Iyer et al. (2022). It contained residual protein (53.9 mg·g^−1^ dry mass) and polysaccharides / sugars (glucose equivalent of 88.9 mg·g^−1^ dry mass). Given that the plant mass was previously treated with cellulase as part of the protein extraction process, the carbohydrate profile of the PDF was expected to comprise of low molecular weight polysaccharides and glucose. Other monomers such as xylose, fucose, and glucuronic acid were expected to remain largely absent as pectinase, xylanase and other cell-wall digesting enzymes were not employed in the process.

The PDF was used at a final concentration of 25 g·L^−1^. This meant that GEM contained about 1.35 g·L^−1^ of proteins and 2.3 g·L^−1^ of carbohydrates; values comparable to CDM.

### Bacterial strains

Four bacterial strains were chosen for B_12_ production, namely, *P. freudenreichii* DSM 4902, *A. caccae* DSM 14662, *A. hallii* DSM 3353, and *R. faecis* DSM 16840, based on the work previously described by Soto-Martín et al. (2020). *A. caccae* DSM 14662, *A. hallii* DSM 3353, and *R. faecis* DSM 16840 were available from the in-house culture stock. *P. freudenreichii* DSM 4902 was purchased from NCIMB (Aberdeen, Scotland, NCIMB 5959).

### Culture revival and growth

Microbial species were revived from frozen stocks using M2GSC (5 mL) in Hungate tubes (sealable with rubber caps) in anaerobic conditions and incubated at 37 °C for five days. The strains were sub-cultured by inoculating 0.5 mL of the culture in 4 mL of fresh M2GSC and incubation for two days. Growth was measured with optical density at 650 nm using Novaspec-II visible spectrophotometer (Pharmacia LKB Biotechnology AB, Uppsala, Sweden).

### Production of B_12_ samples

From the revived stock, 200 μL was inoculated in triplicate in M2GSC, CDM or GEM (10 mL) and gently mixed through repeated aspiration. The culture was statically incubated at 37 °C for 48 h. Sample (1 mL) was drawn at time points 0 h, 6 h, 12 h, 24 h, 30 h, 36 h and 48 h and centrifuged at 10 000 × *g* for 20 min at 4 °C (Biofuge (Fresco), Heraeus Instruments, Germany). The supernatant was recovered and analysed for B_12_ content. The pellet was resuspended in PBS buffer (1 mL) and slowly cooled to −80 °C by first placing them in an ice-bath during sample collection, followed by incubation in −20 °C freezer overnight. Samples were then placed in a −80 °C freezer for a couple of days. This was performed to facilitate gradual growth of ice crystals to pierce and rupture the bacterial cell wall. Samples were then thawed to 4 °C overnight and sonicated (MSE Soniprep 150, UK) in two 10 s bursts to rupture the cells. The cells were centrifuged at 10 000 × *g* for 20 min and the supernatant was recovered and analysed for B_12_ content.

### B_12_ Quantification using LC-MS

B_12_ estimation was performed using mass spectrophotometry in tandem with liquid chromatography as described elsewhere (Jiang et al., 2003). Calibration standards (all from Sigma-Aldrich; Saint Louis, USA) used were adenosylcobalamin, hydroxycobalamin, and methylcobalamin at concentrations 0.3 ng·mL^−1^, 1.0 ng·mL^−1^, 3.0 ng·mL^−1^, 10 ng·mL^−1^, 30 ng·mL^−1^, 100 ng·mL^−1^ and 300 ng·mL^−1^ in dimethyl sulphoxide (DMSO). Cyanocobalamin (internal standard) was used at 2 μg·mL^−1^. Samples or standards were injected at 100 μL with 10 μL internal standard. Details of the LC-MS setup is provided in **S3 Text**.

### Literature review

To retrieve previous literature on B_12_ production in *P. freudenreichii*, the search structure:

~~~
(propionibacterium AND (cobalamin OR B12))
~~~

was used in PubMed and Scopus. The relevant titles were selected and analysed by eye to obtain values for B_12_ yield, the carbohydrate concentration used (Carbon g·L^−1^), fermentation time (Time h), cobalt salt used (mg·L^−1^) and added DMB (mg·L^−1^). These values were used for the construction of a univariate-scaled PCA model to help compare and contextualise the B_12_ yields obtained in the work described herein. The details of the study and B_12_ are provided in **S1 Table**.

### Statistical Analysis

All statistical analyses were performed using R (4.2.2) (R Core Team, 2021) using the tidyverse (2.0.0) (Wickham et al., 2019) package for data processing and visualisation. Normality of the data distribution was checked using the Shapiro-Wilks test. The Carbon (g·L^−1^), Time (h), and B_12_ (mg·L^−1^) values were log_10_ transformed to achieve normality.

The k-means analysis followed by univariate-scaled principal component analysis was performed using the cluster (2.1.4) (Maechler et al., 2022), and factoextra (1.0.7) (Kassambara & Mundt, 2020) packages. A dimension reduction model was made using previously published B_12_ production with *P. freudenreichii*. A k-means analysis (MacQueen algorithm) was performed to observe clustering in the raw data. The raw data was then subjected to a univariate-scaled PCA analysis and the clusters established from k-means analysis was overlaid on the plot to observe their relative positions. This PCA model was then used to interpolate the relative position of the data points representing the experimental conditions described in our study. This was used to understand which cluster of experimental conditions our setup matched and compare the corresponding B_12_ yields. The raw data and codes can be found at https://doi.org/10.17605/osf.io/3yb2r to reproduce the analyses.

## Results and discussion

### Growth profiles

Four bacterial strains, namely, *A. caccae* DSM 14662, *A. hallii* DSM 3353, *R. faecis* DSM 16840 and *P. freudenreichii* DSM 4902 were incubated in CDM, GEM and M2GSC over a period of 48 hours. The growth profiles are provided in Figure 2.

**Figure 2.**
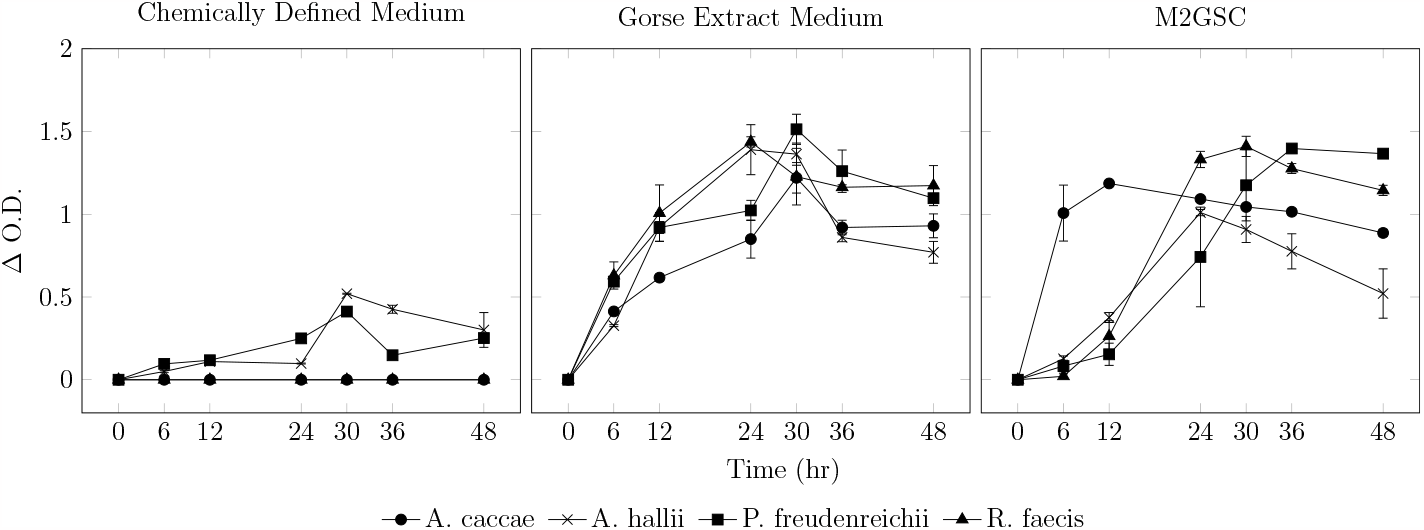
Growth profile of *A. caccae* DSM 14662, *A. hallii* DSM 3353, *P. freudenreichii* DSM 4902, and *R. faecis* DSM 16840 in chemically defined medium (CDM), gorse extract medium (GEM) and M2GSC.

All tested strains demonstrated robust growth in M2GSC and GEM media, reaching ΔO.D values >1.0. In contrast, CDM medium only supported the growth of *A. hallii* DSM 3353 and *P. freudenreichii* DSM 4902, with ΔO.D values <0.51.

In previous experiments by Soto-Martín et al. (2020), *A. caccae* DSM 14662 and *R. faecis* DSM 16840 were grown in CDM variants to assess trophic capabilities. They were initially grown in a semi-defined, vitamin-controlled medium using vitamin-free casein acid hydrolysate fortified with L-tryptophan, L-serine, L-threonine, L-glutamine, and L-asparagine to check for auxotrophy for each individual vitamin. Subsequently, they were grown in an amino-acid controlled CDM supplemented with all vitamins to check for auxotrophy for each amino acid. All three strains used here were able to grow well in both versions of CDM used by Soto-Martín et al. (2020). The growth profile in the panel for CDM in Figure 2 differs from previous reports, and may be explained by the pre-culture conditions. Soto-Martín et al. (2020) subjected the microbes through two passages in CDM before measuring their growth, while in this study, the microbes were revived in M2GSC and directly inoculated in CDM.

The GEM and CDM used were identical in their base compositions of added vitamins (as well as absence of natural cobalamins), minerals, micronutrients, and precursors. The key difference lay in their carbohydrate and protein sources. While GEM contained residual proteins and peptides, CDM contained purified amino acids. This difference may have contributed to the lack of growth by *A. caccae* DSM 14662 and *R. faecis* DSM 16840 in CDM. Certain bacterial species are known to prefer peptides as a nitrogen source rather than individual amino acids (Kampen, 2014) under anaerobic conditions (Cotta & Russell, 1982). The uptake of individual amino acids is energetically demanding and requires active Na^2+^ mediated transmembrane transport (Heyne et al., 1991) and is the primary rate-limiting step in nitrogen uptake (Cotta & Russell, 1982). *A. caccae* DSM 14662 and *R. faecis* DSM 16840 are typically cultured in YCFA medium (La Rosa et al., 2019; Marquet et al., 2009) and the cobalamin-devoid, free-amino-acid based CDM may have been energetically incompatible for their growth.

### B_12_ production in experimental species

The extent of vitamin B_12_ production in the experimental species is shown in Table 1 below.

**Table 1:** Total B_12_ measured in experimental fractions at 48 h incubation

| Species | Media | Lysate | Supernatant |
| --- | --- | --- | --- |
| *A. caccae* | GEM | 16.10±7.59 | N/D |
|  | CDM | N/G | N/G |
|  | M2GSC | N/D | N/D |
| *A. hallii* | GEM | 7.40±5.90 | N/D |
|  | CDM | N/D | N/D |
|  | M2GSC | N/D | N/D |
| *P. freudenreichii* | GEM | 15.21±1.00 | N/D |
|  | CDM | N/D | N/D |
|  | M2GSC | 4.23±0.60<sup>a</sup> | 7.95±1.40 |
| *R. faecis* | GEM | 16.34±0.92 | N/D |
|  | CDM | N/G | N/G |
|  | M2GSC | 0.77±0.12<sup>b</sup> | N/D |

Values expressed in mean±standard deviation µg·L<sup>-1</sup>.
Marked values were significantly different from other yields measured across the microbial species for a given medium.
N/D = below detection level, N/G = no growth
B<sub>12</sub> in all media at time 0 was N/D.

Vitamin B_12_ was below detection levels at the 0 h time point across all tested media. *P. freudenreichii* DSM 4902 and *R. faecis* DSM 16840 were able to produce B_12_ at the 48 h end-point in M2GSC as shown in Table 1. *P. freudenreichii* DSM 4902 showed B_12_ in the supernatant as well as the cell lysate fraction.

Vitamin B_12_ was detected in the lysates of *A. caccae* DSM 14662 and *A. hallii* DSM 3353 cultured in GEM. In contrast, no B_12_ was observed in M2GSC, despite it being the ideal growth medium. DMB was exclusively added to GEM and was likely critical to B_12_ production. The growth of *A. caccae* and *A. hallii* in M2GSC without detectable B_12_ in the lysates could be attributed to the synthesis of pseudo-B_12_ vitamers, that were not detectable by the LC/MS method employed in the study. This may also explain the lower B_12_ levels in M2GSC compared to GEM for *P. freudenreichii* DSM 4902 and *R. faecis* DSM 16840. Although these microbes demonstrated the ability to synthesize biologically active vitamers in anaerobic conditions, the addition of DMB likely alleviated the metabolic burden required for *de novo* synthesis, resulting in an increased total B_12_ production in GEM compared to M2GSC.

In this experiment, B_12_ measurement was performed using LC/MS, capable of detection at ng quantities in the sample. Physiologically, such quantities are far higher than the metabolic requirements of the cell (Burgess et al., 2009). This suggests that the chosen bacterial species were able to overproduce B_12_, but the experimental conditions, or perhaps the media design was unsuitable for any real commercial application. Moreover, this contributes towards confirmation of the speculation (Engels et al., 2016) that *A. hallii* (formerly *E. hallii*) could produce B_12_ vitamers in the gut.

Concerning probiotic applications, effective vitamin production necessitates high production in the extracellular environment. Since the chosen microbes failed to meet both requirements, they may not be ideal candidates for probiotic applications.

Of particular concern was the observed low B_12_ yields for *P. freudenreichii* DSM 4902, which served as the positive control for this experiment. Generally, *P. freudenreichii* DSM 4902 has yields *≥*0.2 μg·mL^−1^, yet the corresponding values noted in Table 1 were about 13*×* lower. A comparative analysis with other literature investigating *P. freudenreichii* for B_12_ production was conducted to explore potential causes.

### Component analyses using previous literature

The systematic online searches yielded 1651 hits from which 230 research titles were retrieved post title screening. A final list of 36 titles were selected based on the relevance of the research described therein. The data points for the concentration of the carbon source (g·L^−1^), cobalt (mg·L^−1^), DMB (mg·L^−1^) and total fermentation time (hours) were retrieved and is presented in **S1 Table**. This only included cases of batch culturing of *P. freudenreichii*.

The data in **S1 Table** was first subjected to k-means clustering to look for grouping patterns in the various experimental conditions that may correspond to B_12_ yields. Based on the “elbow method”, three groups were deemed ideal to explain clustering in the data (**S Codes**). Next, the data was subjected to a univariate-scaled PCA analysis and the corresponding grouping schema was overlaid to visualise their clustering on the PCA plot, as shown in Figure 3.

**Figure 3.**
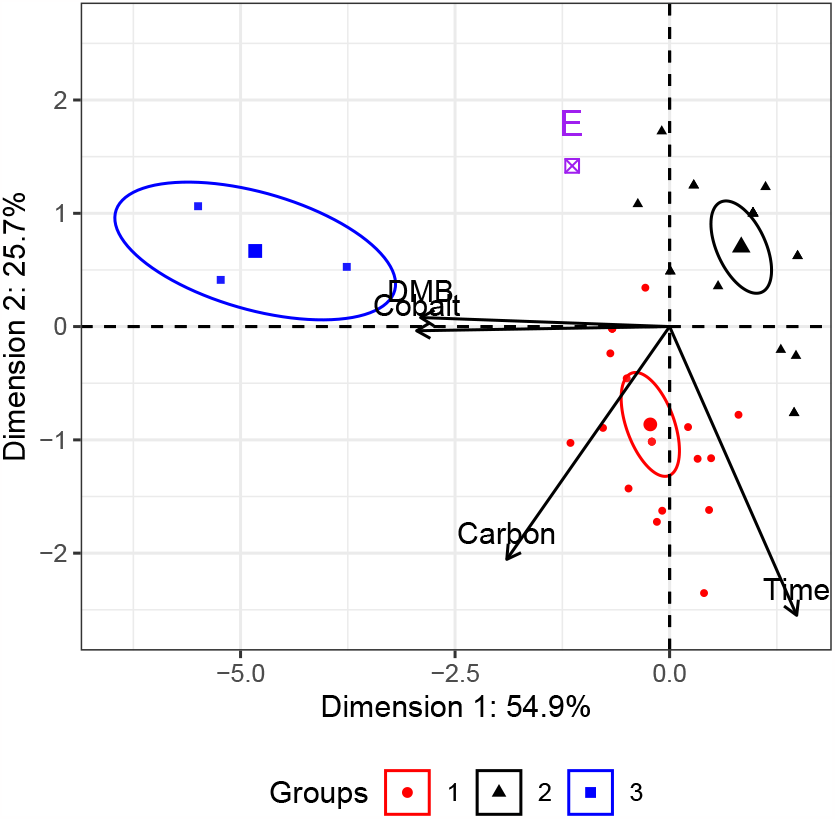
Univariate-scale PCA plot of the experimental conditions employed for B_12_ production using *P. freudenreichii* in previous publications. The point marked ‘E’ refers to the condition employed in this experiment. The position of the ‘E’ point was predicted by the model, which appears to associate to cluster 2. Ellipses represent 95% confidence limits of each cluster.

In Figure 3, the PCA model could account for 80.6% of the total observed variance in the data across Dimensions 1 and 2. The factors in the model, namely Carbon, Cobalt, DMB and Time accounted for 15.7%, 37.9%, 36.8% and 9.6% of the variance accounted in Dimension 1. In Dimension 2, Carbon, Cobalt, DMB and Time accounted for 39.4%, >0.1%, 0.1% and 60.6% of the variance.

Cluster 1 represents conditions with media employing high concentrations of carbon source, as well as a longer fermentation time, Cluster 2 represents the absence of precursors and low carbon source concentrations, and Cluster 3 represents high use of the precursors.

The point labelled “E” represents the experimental conditions and plots the closest to the mid-point of Cluster 2 (*d* = 0.718) compared to Clusters 1 (*d* = 2.57) and 3 (*d* = 5.86), where *d* represents the Euclidean distance between the points. This implies that the experimental condition described in this work was similar to that represented by Cluster 2. The median value of the conditions represented in each cluster is presented in Table 2.

**Table 2:** Median values of the experimental parameters and B_12_ yield represented in the clusters identified using k-means and PCA analysis.

| Groups | Carbon | Cobalt | DMB | Time | B<sub>12</sub> |
| --- | --- | --- | --- | --- | --- |
| Cluster 1 | 30 (25-35) | 5.5 (5-12.7) | 10 (0.9-15.0) | 120 (72-16.0) | 5.7 (1.3-16.1) |
| Cluster 2 | 10 (8.4-10)<sup>a</sup> | 0 (0-0) | 0 (0-0) | 72 (72-102) | 0.22 (0.16-2.21) |
| Cluster 3 | 40 (31.3-42.0) | 40 (29-45) | 70 (70-70) | 48<sup>b</sup> (47.5-48.0) | 6.7 (4.2-7.4) |
| Experimental | 6<sup>a</sup> | 10 | 20 | 48<sup>b</sup> | 0.01 |

Values expressed as median (25% quantile to 75% quantile) to indicate the data distribution.
Carbon represents carbohydrates in the media (g·L<sup>-1</sup>).
Cobalt and DMB (5,6-dimethylbenzimidazole) are expressed in mg·L<sup>-1</sup>.
Time represents the total fermentation time (h).
B<sub>12</sub> was the final reported yield (mg·L<sup>-1</sup>).
Values with the same superscript were statistically similar using one-sample t-test.

The yield in Cluster 2 (0.2 mg·L^−1^) was significantly lower than in Clusters 1 (5.7 mg·L^−1^) and 3 (6.7 mg·L^−1^), respectively (F_(2, 44)_ = 5.298, p = 0.009). These findings suggest that the availability of carbon source (via carbohydrates), was lower in the experimental setup described here compared to previous publications. As presented in Table 1, B_12_ content was predominantly retained within the cell lysate, indicating that the microbes did not secrete B_12_ extracellularly. Consequently, to achieve desired B_12_ yields, it is imperative to supply essential nutrients and precursors at sufficiently high concentrations, as indicated by the values presented in Table 2. **S1 Fig** illustrates boxplots comparing the levels of Cobalt, Carbon, DMB, and Time for each cluster.

Thus the data from Table 2 suggest that the yields of B_12_ are impacted when the carbohydrate concentration in the media is >30 g·L^−1^.

## Limitations and scope for further work

This experiment was designed to investigate two primary objectives: first, to determine the growth potential of selected gut microbes on media composed of PDF generated from Gorse wastes, and second, to assess their ability to produce vitamin B_12_. The focus of this study was primarily to test B_12_ production in the candidate microbes and their ability to grow on Gorse waste media, rather than delve into the intricate mechanisms behind vitamin B_12_ production or enhancing yields for industrial applications.

Consequently, there were some aspects of this experiment that may warrant additional investigation. For example, *A. hallii* has previously been reported to produce pseudo-vitamin B_12_ (Belzer et al., 2017). Thus, while this experiment could demonstrate that biologically active B_12_ was produced by *A. hallii*, it is difficult to conclude if the microbe can facultatively produce the bioactive vitamer without a control medium devoid of DMB.

Expanding on the investigation, it’s worth noting that other colonic microbes, such as *Lactobacillus reuteri*, are known to produce pseudo-cobalamin under anaerobic conditions. These microbes present intriguing candidates for future studies exploring their potential to produce the bioactive vitamer when DMB is introduced into the media.

Lastly, the individual vitamers and pseudo-vitamers were not characterised. More work will be required to understand if there are unique stoichiometries in which these vitamers are produced.

## Conclusion

The investigation could show for the first time that *A. hallii, R. faecis* and *A. caccae* can produce vitamin B_12_, albeit in very small quantities. However, these candidates are not suitable candidates for commercial B_12_ production. The Protein Depleted Fraction (PDF) stands out as a valuable carbon and nitrogen source, capable of supporting the growth of gut microbes. This sets the stage for developing circular processes to cultivate microbes with higher B_12_ production potential and pave the way for sustainable probiotic products.

## Supporting information

Supplementary_Codes

## Abbreviations

CDM: chemically defined medium
GEM: Gorse extract medium
PDF: protein depleted fraction
PCA: principal component analysis

## Supporting information

### M2GSC recipe

The recipe for the M2GSC medium used for the growth of the experimental cells is provided section 1.1 in the Supplementary Materials file.

### Chemically defined medium recipe

The recipe for the the chemically defined medium (CDM) used for the growth of the experimental cells is provided section 1.2 in the Supplementary Materials file.

### LC/MS conditions

Details of the LC/MS setup to measure B_12_ is provided section 1.3 in the Supplementary Materials file.

### S Codes

The codes used to analyse the data is annotated and available the Supplementary Codes file.

### S1 Fig. Measures of experimental parameters in k-means clusters

Median levels of B_12_ yield, carbon source, precursors, and fermentation time in the k-means groups, shown in Figure 3. This can be found in the Supplementary Materials file.

### Supplementary text and details

Details on media recipe and LC/MS conditions. This can be found in the Supplementary Materials file.

### S1 Table. B_12_ yields in previous publications

This can be found in the Supplementary Materials file.

## Acknowledgments

We thank Freda Farquharson and Jenny Martin for their help and support in the laboratory. We thank Wiley Barton for their comments on the manuscript. This work was funded by the Scottish Government through Rural and Environment Science and Analytical Services Division (RESAS) as part of its strategic research programme. This work was performed as part of A.I’s PhD project (2017 to 2021).

---

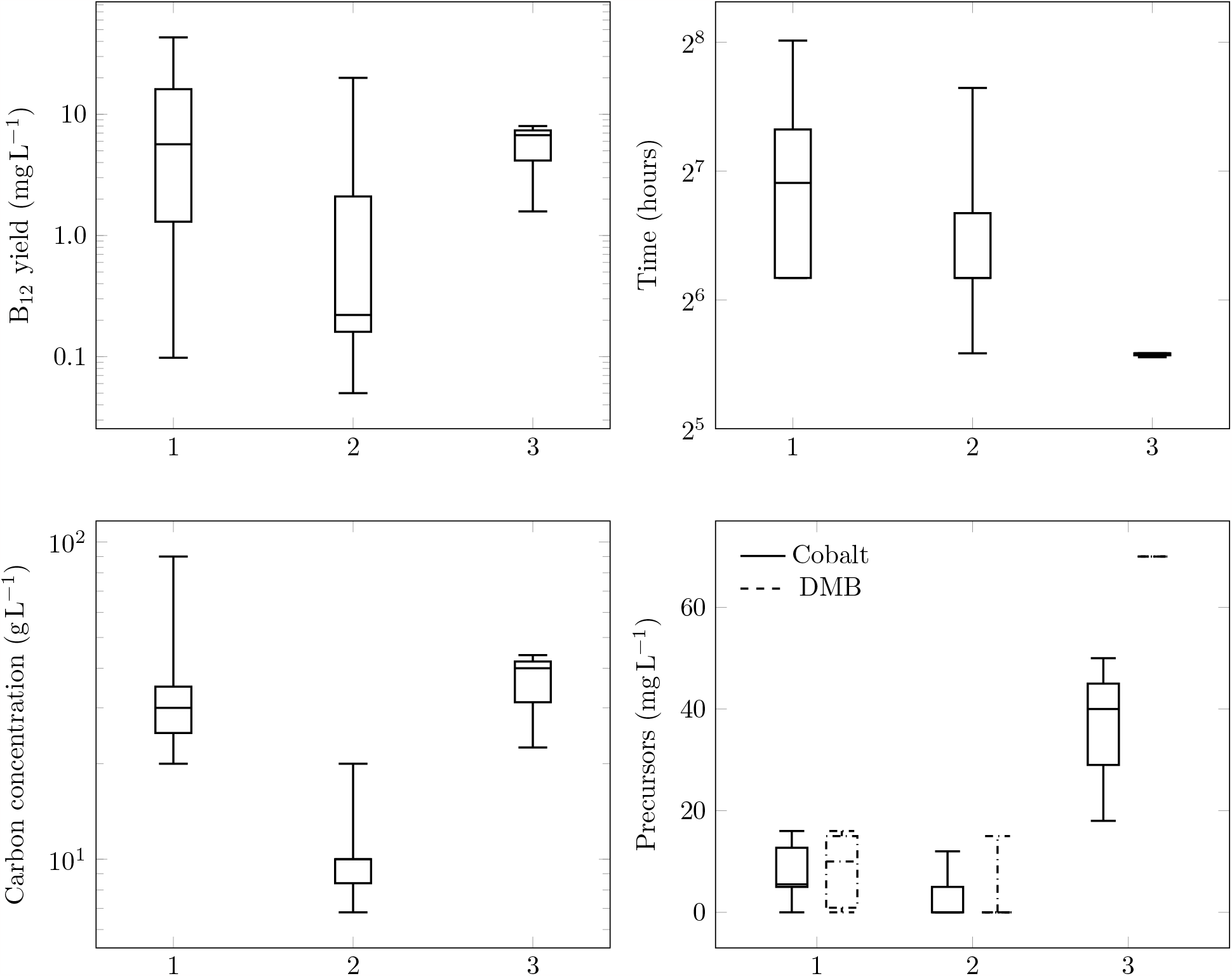

## Notes

https://osf.io/3yb2r/

### Competing Interest Statement

The authors have declared no competing interest.

### Summary of Updates

1. The abstract is changed to provide more details on the methodology. 2. The Introduction has been expanded to provide more background on the topic. The context and research interest driving this investigation is elaborated further. 3. Some paragraphs in the results and discussion has been reworded to express the idea more clearly. 4. The conclusion section has been streamlined, and now only highlights the major outcomes of the study. Limitations of the study remains a separate section and irrelevant texts have now been removed. 5. The layout of the manuscript has also been changed and the citation style is now APA.

