## Supplementary_Materials for "Gorse (*Ulex europeaus*) wastes with 5,6-dimethyl benzimidazole supplementation can support growth of vitamin B_12_ producing commensal gut microbes"

### Supplementary Material for: Vitamin $B_{12}$ synthesis in three commensal gut microbes and their application in valorising Gorse (*Ulex europeaus*) wastes.

Ajay Iyer[1,3], Eva C. Soto-Martin[2], Gary Cameron[1], Charles S. Bestwick[1], Sylvia H. Duncan[1], and Wendy R. Russell*[1]

[1]University of Aberdeen, Rowett Institute, Aberdeen AB25 2ZD, Scotland
[2]University of Aberdeen, IMS, Aberdeen AB25 2ZD, Scotland
[3]Teagasc, Fermoy, Co Cork, Ireland.

#### 1 Supplementary texts

Additional details and codes may be found in the OSF repository: https://osf.io/3yb2r/

##### 1.1 M2GSC recipe

Bacto™ Casitone (10 g·L$^{-1}$), Bacto™ Yeast Extract (2.5 g·L$^{-1}$), sodium bicarbonate ($NaHCO_3$, 4.0 g·L$^{-1}$), glucose (2 g·L$^{-1}$), soluble starch (2 g·L$^{-1}$), cellobiose (2 g·L$^{-1}$), clarified rumen fluid (100 mL·L$^{-1}$), cysteine HCl (1 g·L$^{-1}$), dipotassium hydrogen phosphate ($K_2HPO_4$, 0.4 g·L$^{-1}$), potassium dihydrogen phosphate ($K_2HPO_4$, 0.4 g·L$^{-1}$), ammonium sulphate ($(NH_4)_2SO_4$, 0.8 g·L$^{-1}$), sodium chloride (NaCl, 0.8 g·L$^{-1}$), magnesium sulphate ($MgSO_4$, 80 mg·L$^{-1}$), calcium chloride ($CaCl_2$, 80 mg·L$^{-1}$) and resazurin (1 µg·L$^{-1}$).

##### 1.2 Chemically defined medium (CDM)

**Nitrogen sources (in levo-chirality):** alanine (6.2 mg·L$^{-1}$), arginine (7.3 mg·L$^{-1}$), aspartate (4.1 mg·L$^{-1}$), aspartic acid (5.4 mg·L$^{-1}$), glutamine (4.2 mg·L$^{-1}$), glutamic acid (7.1 mg·L$^{-1}$), glycine (4.1 mg·L$^{-1}$), histidine (2.5 mg·L$^{-1}$), isoleucine (6.7 mg·L$^{-1}$), leucine (10.4 mg·L$^{-1}$), lysine (6.4 mg·L$^{-1}$), methionine (2.9 mg·L$^{-1}$), phenylalanine (5.3 mg·L$^{-1}$), proline (3.8 mg·L$^{-1}$), serine (4.9 mg·L$^{-1}$), threonine (4.8 mg·L$^{-1}$), tryptophan (1.9 mg·L$^{-1}$), valine (6.3 mg·L$^{-1}$), cysteine (10 mg·L$^{-1}$).

*

**Carbon sources (in dextro-chirality):** glucose ($2\,g{\cdot}L^{-1}$), galactose ($2\,g{\cdot}L^{-1}$), cellobiose ($2\,g{\cdot}L^{-1}$) and potassium hydrogen carbonate ($KHCO_3$, $40\,mg{\cdot}L^{-1}$).

**Mineral sources:** dipotassium hydrogen phosphate ($K_2HPO_4$, $0.2\,mg{\cdot}L^{-1}$), ammonium sulphate ($(NH_4)_2SO_4$, $4\,mg{\cdot}L^{-1}$), sodium chloride (NaCl, $4\,mg{\cdot}L^{-1}$), magnesium sulphate, heptahydrate ($MgSO_4 \cdot 8\,H_2O$, $0.4\,mg{\cdot}L^{-1}$), calcium chloride ($CaCl_2$, $0.4\,mg{\cdot}L^{-1}$) and potassium dihydrogen phosphate ($KH_2PO_4$, $2\,mg{\cdot}L^{-1}$), ferrous sulphate heptahydrate ($FeSO_4 \cdot 7\,H_2O$, $21\,mg{\cdot}L^{-1}$), zinc sulphate heptahydrate ($ZnSO_4 \cdot 7\,H_2O$, $1.8\,mg{\cdot}L^{-1}$), boric acid ($H_3BO_3$, $5\,mg{\cdot}L^{-1}$), cobalt chloride hexahydrate ($CoCl_2 \cdot 6\,H_2O$, $10\,mg{\cdot}L^{-1}$), nickel chloride hexahydrate ($NiCl_2 \cdot 6\,H_2O$, $0.1\,mg{\cdot}L^{-1}$), copper chloride dihydrate ($CuCl_2 \cdot 2\,H_2O$, $0.1\,mg{\cdot}L^{-1}$), manganese chloride tetrahydrate ($MnCl_2 \cdot 4\,H_2O$, $0.7\,mg{\cdot}L^{-1}$), sodium molybdate dihydrate ($Na_2MoO_4 \cdot 2\,H_2O$, $0.5\,mg{\cdot}L^{-1}$), sodium selenite pentahydrate ($Na_2SeO_3 \cdot 5\,H_2O$, $0.1\,mg{\cdot}L^{-1}$), sodium tungstate dihydrate ($Na_2WO_4 \cdot 2\,H_2O$, $0.1\,mg{\cdot}L^{-1}$).

**Others:** 5,6-dimethylbenzimidazole (DMB, $20\,mg{\cdot}L^{-1}$), pyridoxine HCl ($1\,ng{\cdot}L^{-1}$), pyridoxal HCl ($1\,ng{\cdot}L^{-1}$), pyridoxamine HCl ($1\,ng{\cdot}L^{-1}$), thiamine HCl ($0.5\,ng{\cdot}L^{-1}$), Ca-D-pantothenate (B5, $0.5\,ng{\cdot}L^{-1}$), pantetheine ($0.5\,ng{\cdot}L^{-1}$), nicotinic acid ($0.5\,ng{\cdot}L^{-1}$), nicotinamide ($0.5\,ng{\cdot}L^{-1}$), riboflavin ($0.5\,ng{\cdot}L^{-1}$), folic acid ($0.5\,ng{\cdot}L^{-1}$), p-amino benzoic acid ($0.5\,ng{\cdot}L^{-1}$), menadione ($0.1\,ng{\cdot}L^{-1}$), lipoic acid ($0.5\,ng{\cdot}L^{-1}$), pimelic acid ($0.2\,ng{\cdot}L^{-1}$), biotin ($0.2\,ng{\cdot}L^{-1}$).

**Nucleotides:** adenine ($10\,mg{\cdot}L^{-1}$), cytosine ($10\,mg{\cdot}L^{-1}$), thymine ($10\,mg{\cdot}L^{-1}$), xanthine ($10\,mg{\cdot}L^{-1}$), guanine ($10\,mg{\cdot}L^{-1}$), uracil ($10\,mg{\cdot}L^{-1}$), orotic acid ($5\,mg{\cdot}L^{-1}$) and thymidine ($10\,mg{\cdot}L^{-1}$).

#### 1.3 LC/MS conditions for $B_{12}$ estimation

Liquid chromatography was performed using An ACE Excel 3 µ $C_{18}$ (150 mm × 2.1 mm) column maintained at 50 °C. Elution was performed across a gradient using solvent A (0.1% $v{\cdot}v^{-1}$ formic acid) and solvent B (methanol). The complete programme lasted 10 min at a flow rate of $200\,\mu L{\cdot}min^{-1}$ with a sample/standard injection volume of 5 µL. The column was maintained at 90:10:: Solvent A: Solvent B which was rapidly raised to 0:100::Solvent A:Solvent B at 4 min and maintained for 2 min. This was followed by a rapid drop to 90:10::Solvent A: Solvent B and maintained until the final 10 min runtime.

The mass spectrophotometry setup was performed using positive ion electrospray at 4 kV. Capillary pressure was 375 °C and collision pressure was 1.8 mTorr. Q1/Q3 peak width was 0.7. Approximate retention times (min) were: hydroxycobalamin (5.75), methyl-cobalamin (6.95), cyanocobalamin (6.25) and adenosylcobalamin (6.61). Calibration was performed using a quadratic transformation with $\frac{1}{x^2}$ weighting.

#### 2 Supplementary figure

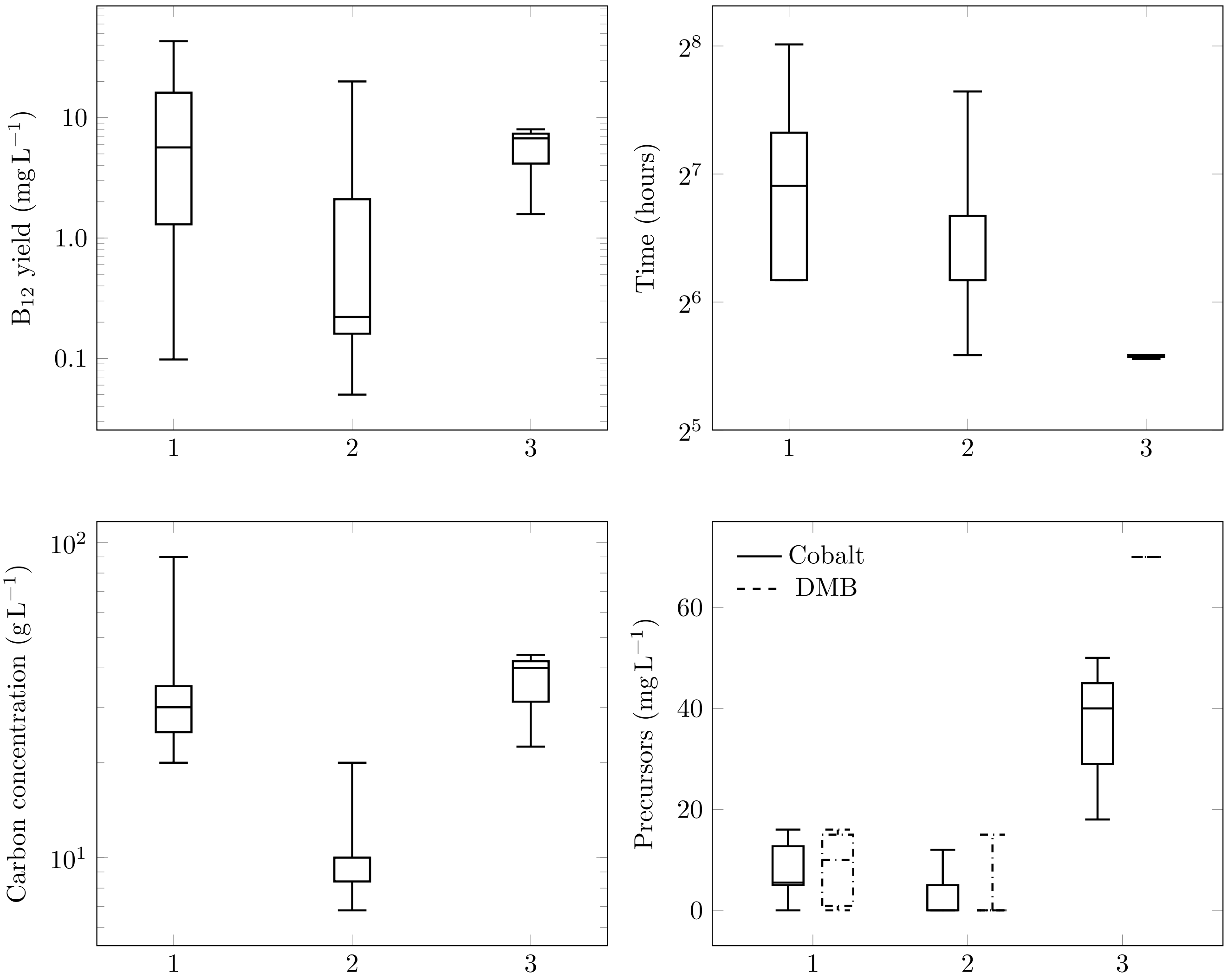


Supplementary Figure 1: Median levels of $B_{12}$ yield, carbon source, precursors, and fermentation time in the k-means groups.

### 3 Supplementary table

Table 1: $B_{12}$ production in previous literature. Data used for PCA analysis shown in Figure 3 in main document.

| Medium | DMB | Cobalt | $B_{12}$ | Carbon | Time | Ref |
|---|---|---|---|---|---|---|
| Casio peptone medium | 70 | 18 | 6.73 | 40 | 48 | [1] |
| Cereal matrix medium | 15 | 5 | 1.5 | 20 | 144 | [2] |
| Cheese propionic acid medium | 0 | 0 | 0.124 | 6.8 | 108 | [3] |
| Cheese whey + yeast extract | 0 | 0 | 0.12 | 6.8 | 108 | [3] |
| Cheese whey + yeast extract | 5 | 5 | 20 | 7 | 168 | [4, 5, 6] |
| Cornsteep liquor | 0.9 | 16 | 12.5 | 41 | 132 | [7] |
| Cornsteep liquor | 0.9 | 12.7 | 21.5 | 56 | 84 | [8] |
| Cornsteep liquor | 0.9 | 5 | 18.2 | 35 | 160 | [9] |
| Cornsteep liquor | 0.9 | 12.7 | 35.2 | 20 | 72 | [10] |
| Cornsteep liquor | 0.9 | 12.7 | 43.5 | 30 | 258 | [11] |
| Cornsteep liquor | 14.6 | 5 | 2.06 | 90 | 72 | [12] |
| Lactate Media | 0 | 5 | 0.05 | 9 | 200 | [13] |
| Lactate Media | 0 | 5 | 0.16 | 9 | 200 | [13] |
| Lactate Media | 0 | 0 | 0.161 | 10 | 72 | [14] |
| Lactate Media | 0 | 0 | 0.227 | 10 | 72 | [14] |
| Lactate Media | 0 | 0 | 0.215 | 10 | 72 | [14] |
| Lactate Media | 0 | 0 | 0.681 | 10 | 72 | [14] |
| Lactate Media | 0 | 0 | 0.188 | 10 | 72 | [14] |
| Lactate Media | 0 | 0 | 0.168 | 10 | 72 | [14] |
| Lactate Media | 0 | 0 | 0.211 | 10 | 72 | [14] |
| Lactate Media | 0 | 0 | 0.221 | 10 | 72 | [14] |
| pABA Medium | 15 | 5 | 0.625 | 20 | 168 | [15] |
| Peptone Medium | 15 | 10 | 5.53 | 10 | 96 | [16] |
| Peptone Medium | 15 | 10 | 3.81 | 10 | 96 | [16] |
| Peptone Medium | 16 | 2 | 3.039 | 20 | 168 | [17] |
| Soya Medium | 15 | 15 | 0.528 | 20 | 100 | [18] |
| Soya Medium | 70 | 40 | 1.58 | 44 | 48 | [19] |
| Spent media | 0 | 0 | 0.95 | 10 | 72 | [20] |
| Sunflower medium | 15 | 5.5 | 1.6 | 40 | 160 | [21] |
| Tofu waste | 0 | 12 | 3.2 | 10 | 48 | [22] |
| Whey permeate media | 0 | 0 | 2.5 | 7.8 | 72 | [23] |
| yeast extract | 0 | 0 | 0.06 | 20 | 72 | [24] |
| yeast extract | 0 | 0 | 1.68 | 10 | 144 | [25] |
| yeast extract | 0 | 10 | 0.087 | 20 | 48 | [26] |
| yeast extract | 0 | 0 | 13.9 | 25 | 120 | [27] |
| yeast extract | 0.9 | 12.7 | 42.5 | 35 | 84 | [28] |
| yeast extract | 70 | 50 | 8 | 22.5 | 47 | [29] |
| yeast extract | 14.6 | 5 | 2.6 | 7.8 | 72 | [30] |
| yeast extract | 14.6 | 5 | 1.7 | 7.8 | 72 | [30] |
| yeast extract | 15 | 5.5 | 1.3 | 36 | 160 | [21] |
| Yeast extract + lactate | 15 | 5 | 0.223 | 32.8 | 120 | [31] |
| Yeast extract + lactate | 15 | 5 | 0.205 | 32.8 | 120 | [31] |
| Yeast extract + lactate | 15 | 5 | 0.098 | 32.8 | 120 | [31] |
| Yeast extract + lactate | 10 | 10 | 16.13 | 30 | 72 | [32] |
| Yeast extract + lactate | 10 | 10 | 10.93 | 30 | 72 | [32] |
| Yeast extract + lactate | 10 | 10 | 5.66 | 30 | 72 | [32] |
| Yeast extract + lactate | 10 | 10 | 12.06 | 30 | 72 | [32] |

Time expressed in h.
Carbohydrates (carbon) expressed in $g \cdot L^{-1}$.
Cobalt and 5,6-dimethylbenzimidazole (DMB) expressed in $mg \cdot L^{-1}$.
