## Supplementary_Codes for "Gorse (*Ulex europeaus*) wastes with 5,6-dimethyl benzimidazole supplementation can support growth of vitamin B_12_ producing commensal gut microbes"

### Supplementary Material for: Vitamin $B_{12}$ synthesis in three commensal gut microbes and their application in valorising Gorse (*Ulex europeaus*) wastes.

Ajay Iyer[1,3], Eva C. Soto Martín[2], Gary Cameron[1], Petra Louis[1], Sylvia H. Duncan[1], Charles S. Bestwick[1], and Wendy R. Russell[1]

[1]University of Aberdeen, Rowett Institute, Aberdeen, AB25 2ZD, Scotland.
[2]University of Aberdeen, IMS, Aberdeen, AB25 2ZD, Scotland.
[3]Teagasc, Moorepark Food Research Centre, Fermoy, Co Cork, P61 C996, Ireland.

2023-12-18

#### Load the required libraries

These are the packages that will be used across all the analyses described here. Please make sure they are installed in your system.

```r
library(tidyverse) #Set of packages for data wrangling and visualisation.
library(factoextra) #Package for PCA analysis.
library(reshape2) #Package to reshape data structure.
library(cluster) #Package to for k-means.
library(formatR) #Package to make this Markdown render correctly.
```

#### 1 Plotting the species growth curves

In this section, the O.D. values are plotted to obtain the growth curves. The data is already blank subtracted. The output in the main manuscript was plotted using pgfplots in LaTeX for better aesthetics, but the outcome is the same and reproducible. The O.D. values are stored in the file `Table_Supplementary1.csv`

The Supplementary Table 1 is imported as `df1`

```r
df1<-read.csv("Table_Supplementary1.csv")

df1%>%
  group_by(Time, Medium, Species)%>%
  summarise(DOD = mean(OD),
            SD = sd(OD))%>%
  ggplot(aes(x=Time, y=DOD, shape=Species))+
  geom_point()+
  geom_line()+
  geom_errorbar(aes(ymin=DOD-SD, ymax=DOD+SD), width=1.2)+
  facet_grid(~Medium)+
  labs(y = "Delta OD",
```

```
      x = "")+
  theme_bw()+
  theme(legend.position = "bottom",
        strip.background = element_rect(colour = NA, fill = NA))
```

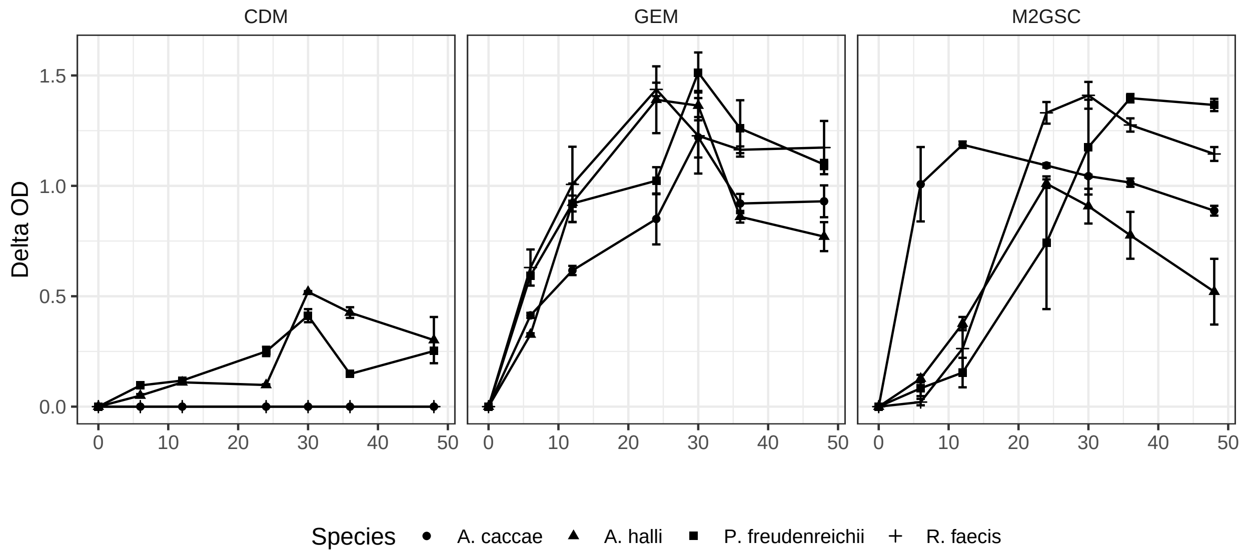


### 2 Analysis of $B_{12}$ yields

In this section, the data table containing the $B_{12}$ yields in for each species, in each medium at the 48 h time point is loaded to run statistical tests. The data is stored in the file `Table_Supplementary2.csv`.

The Supplementary Table 2 is imported as object `df2`.

```
df2<-read.csv("Table_Supplementary2.csv")

df2%>%
  filter(Fraction == "Lysate"
         & Medium == "Gorse")%>%
  aov(formula =  Value ~ Species)%>%
  summary()
```

```
##             Df Sum Sq Mean Sq F value Pr(>F)
## Species      3  163.9   54.65   2.316  0.152
## Residuals    8  188.7   23.59
```

```
df2%>%
  filter(Fraction == "Lysate"
         & Medium == "M2GSC")%>%
  aov(formula =  Value ~ Species)%>%
  summary()
```

```
##             Df Sum Sq Mean Sq F value   Pr(>F)
```

```
## Species      3  36.70  12.233   131.3 3.83e-07 ***
## Residuals    8   0.75   0.093
## ---
## Signif. codes:  0 '***' 0.001 '**' 0.01 '*' 0.05 '.' 0.1 ' ' 1
```

```
df2%>%
  filter(Fraction == "Lysate"
        & Medium == "M2GSC")%>%
  aov(formula =  Value ~ Species)%>%
  TukeyHSD()
```

```
##   Tukey multiple comparisons of means
##     95% family-wise confidence level
##
## Fit: aov(formula = Value ~ Species, data = .)
##
## $Species
##                                     diff         lwr        upr     p adj
## E.hallii-A.caccae         -2.220446e-16 -0.79795061  0.7979506 1.0000000
## P.freudenreichii-A.caccae  4.230000e+00  3.43204939  5.0279506 0.0000007
## R.faecis-A.caccae          7.733333e-01 -0.02461728  1.5712839 0.0574672
## P.freudenreichii-E.hallii  4.230000e+00  3.43204939  5.0279506 0.0000007
## R.faecis-E.hallii          7.733333e-01 -0.02461728  1.5712839 0.0574672
## R.faecis-P.freudenreichii -3.456667e+00 -4.25461728 -2.6587161 0.0000034
```

### 3 Clustering and dimension reduction of *P. freudenreichii* yields reported in literature

```
df<-read.csv("Table_Supplementary3.csv")%>%
  select(6:10)%>%
  transmute(#
    DMB=as.numeric(DMB.mg.L.),
    Cobalt=as.numeric(Cobalt.mg.L.),
    Carbon=log10(as.numeric(Carbon.g.L.)),
    Time=log10(as.numeric(Time..Hr.)),
    B12=log10(as.numeric(B12.mg.L.)))

head(df) #show the first six lines of the table
```

```
##   DMB Cobalt   Carbon     Time        B12
## 1   0      0 1.30103 1.857332 -1.1881570
## 2  15     15 1.30103 2.000000 -0.2773661
## 3  15     10 1.00000 1.982271  0.7427251
## 4  15     10 1.00000 1.982271  0.5809250
## 5  16      2 1.30103 2.225309  0.4827307
## 6   0      0 1.00000 2.158362  0.2253093
```

```
summary(df) #Describe the data in the table
```

```
##        DMB             Cobalt           Carbon            Time      
##  Min.   : 0.00   Min.   : 0.000   Min.   :0.7782   Min.   :1.672  
##  1st Qu.: 0.00   1st Qu.: 0.000   1st Qu.:1.0000   1st Qu.:1.857  
##  Median : 0.90   Median : 5.000   Median :1.3010   Median :1.857  
##  Mean   :10.21   Mean   : 7.392   Mean   :1.2390   Mean   :1.965  
##  3rd Qu.:15.00   3rd Qu.:10.000   3rd Qu.:1.4868   3rd Qu.:2.090  
##  Max.   :70.00   Max.   :50.000   Max.   :1.9542   Max.   :2.412  
##        B12         
##  Min.   :-1.9144  
##  1st Qu.:-0.6788  
##  Median : 0.2014  
##  Mean   : 0.1105  
##  3rd Qu.: 0.8468  
##  Max.   : 1.6385  
```

In the code chunk above, the the first five columns were not selected as they contained textual data regarding the Year of publication, First Author, publication DOI, the bacterial strain, and the growth medium. These were not relevant factors in statistical analyses. The remaining five columns were imported and converted to `numeric` data format. Carbon, Time and $B_{12}$ were $\log_{10}$ transformed while DMB and Cobalt were unchanged as they contained `0` values.

#### 3.1 Data clustering

In this step the data was clustered using k-means analysis. Clustering was performed excluding the $B_{12}$ column. Since $B_{12}$ was the outcome, the conditions were the input factors using which, the clustering and subsequent dimension reduction models would be constructed.

Furthermore, the last line in Supplementary table contained the experimental conditions which will be scaled and plotted in the subsequent principal component analysis.

```
set.seed(1947)
df2<-scale(df[1:47,1:4]) #Scaling the dataset.
fviz_nbclust(df2, kmeans, k.max = 5, method = "wss")+ #Looking for elbow which appears at 3
geom_vline(xintercept = 3, linetype="dashed")
```

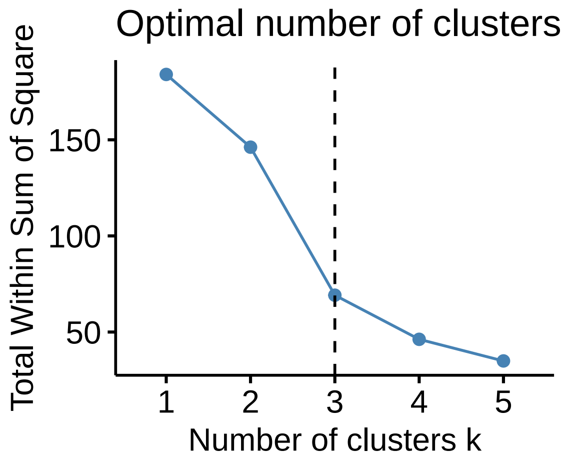


A k-means analysis was performed using the code below. `set.seed()` was used to make the example reproducible.

```r
set.seed(1947)
km <- kmeans(df2, centers = 3, algorithm = "MacQueen",
             nstart = 25, iter.max = 25)

km
```

```
## K-means clustering with 3 clusters of sizes 21, 23, 3
##
## Cluster means:
##           DMB      Cobalt     Carbon       Time
## 1 -0.03906066  0.07817171  0.8407343  0.4295644
## 2 -0.41993352 -0.46129828 -0.8939446 -0.1871562
## 3  3.49291496  2.98941813  0.9684352 -1.5720870
##
## Clustering vector:
##  1  2  3  4  5  6  7  8  9 10 11 12 13 14 15 16 17 18 19 20 21 22 23 24 25 26
##  2  1  2  2  1  2  2  1  1  1  1  1  3  2  2  1  2  2  1  2  2  1  1  2  2  2
## 27 28 29 30 31 32 33 34 35 36 37 38 39 40 41 42 43 44 45 46 47
##  2  2  2  2  2  1  1  1  1  3  1  1  2  2  1  3  2  2  1  1  1
##
## Within cluster sum of squares by cluster:
## [1] 28.678950 28.256931  6.409987
##  (between_SS / total_SS =  65.6 %)
##
## Available components:
##
## [1] "cluster"      "centers"      "totss"        "withinss"     "tot.withinss"
## [6] "betweenss"    "size"         "iter"         "ifault"
```

The clustering groups as described in the output of `km` were appended to the main dataframe `df` so as to assign each row to its corresponding cluster, in the chunk below:

```r
df3<-data.frame(Cluster=as.factor(km$cluster),
                B12=10^(df$B12)[1:47],
                Time=10^(df$Time)[1:47],
                DMB= df$DMB[1:47],
                Cobalt= df$Cobalt[1:47],
                Carbon=10^(df$Carbon)[1:47])
```

Once the clusters were assigned to `df`, the corresponding $B_{12}$ values of the cluster was visualised using boxplots.

To check if the clusters differed significantly in their $B_{12}$ yield, a one-way ANOVA followed by a Tukey *post hoc* test was performed.

```r
df3.aov<-aov(data =df3, B12~Cluster)
summary(df3.aov)
```

```
##             Df Sum Sq Mean Sq F value  Pr(>F)
## Cluster      2   1024   512.0   5.298 0.00868 **
## Residuals   44   4253    96.7
## ---
## Signif. codes:  0 '***' 0.001 '**' 0.01 '*' 0.05 '.' 0.1 ' ' 1
```

```r
TukeyHSD(df3.aov)
```

```
##   Tukey multiple comparisons of means
##     95% family-wise confidence level
##
## Fit: aov(formula = B12 ~ Cluster, data = df3)
##
## $Cluster
##          diff       lwr       upr     p adj
## 2-1 -9.642504 -16.83964 -2.445371 0.0061592
## 3-1 -6.147048 -20.86482  8.570725 0.5726762
## 3-2  3.495456 -11.14211 18.133023 0.8318924
```

The distribution of the data grouped into the k-means clusters was checked using boxplots. In the plot below, the concentrations of $B_{12}$, Cobalt and 5,6-dimethylbenzimidazole in each of the clusters was visualised.

```r
df3%>%
  melt(value.name = "Value", variable.name = "Parameters", id=1)%>%
  group_by(Cluster, Parameters)%>%
  filter(Parameters == "B12" |
           Parameters == "Cobalt" |
           Parameters == "DMB")%>%
  ggplot(aes(x=Cluster, y=Value))+
  geom_boxplot(width=0.4)+
  scale_y_log10()+
  facet_wrap(.~Parameters)+
  ylab("Concentration mg /L")+
  theme_bw()
```

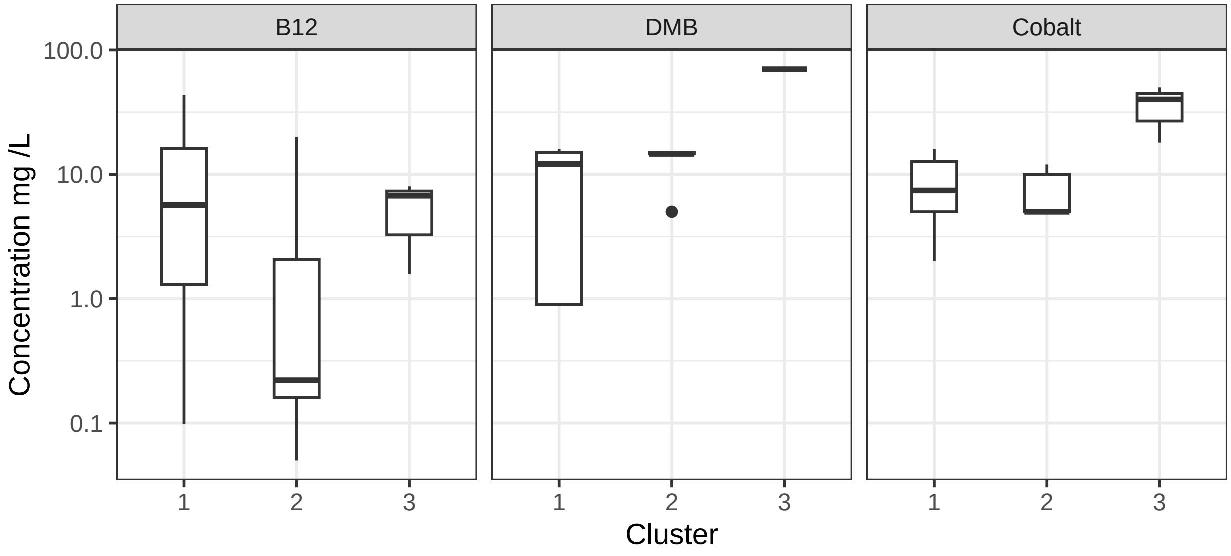


The concentration of the carbon source used in the media is visualised in the figure below.

```r
df3%>%
  melt(value.name = "Value", variable.name = "Parameters", id=1)%>%
  group_by(Cluster, Parameters)%>%
```

```
  filter(Parameters == "Carbon")%>%
  ggplot(aes(x=Cluster, y=Value))+
  geom_boxplot(width=0.4)+
  scale_y_log10()+
  facet_wrap(.~Parameters)+
  ylab("Concentration g /L")+
  theme_bw()
```

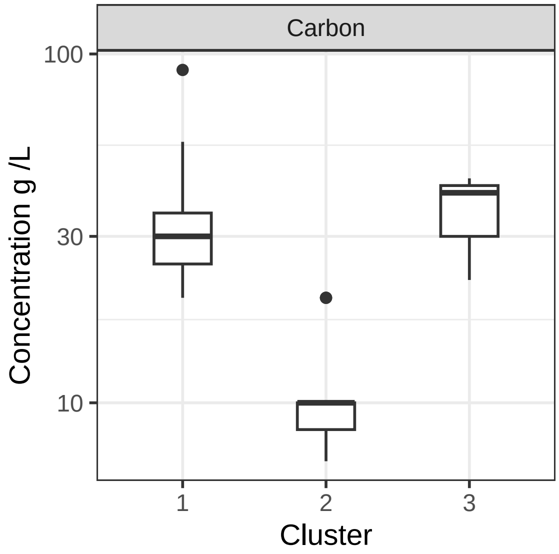


Lastly, the fermentation time used in each cluster is visualized below.

```
df3%>%
  melt(value.name = "Value", variable.name = "Parameters", id=1)%>%
  group_by(Cluster, Parameters)%>%
  filter(Parameters == "Time")%>%
  ggplot(aes(x=Cluster, y=Value))+
  geom_boxplot(width=0.4)+
  scale_y_log10()+
  facet_wrap(.~Parameters)+
  ylab("Time (hr)")+
  theme_bw()
```

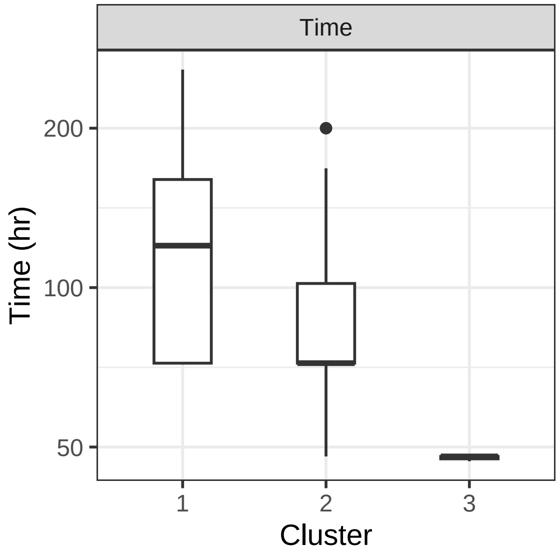


Next, a univariate scaled principal component analysis (PCA) was performed.

```
res.pca<-prcomp(df[1:47,1:4], scale. = TRUE, center = TRUE)

Contrib<-fviz_contrib(res.pca, choice = "var", axes = 1:2)

fviz.plot<-fviz_pca_biplot(res.pca,
                           habillage = as.factor(km$cluster), #Groups based on the k-means clusters.
                           addEllipses = TRUE,
                           col.var = "black",
                           axes = c(1,2),
                           pointsize=1,
                           geom = "point",
                           ellipse.type="confidence", #Ellipses represent confidence limits
                           #mean.point=FALSE,
                           mean.point.size=c(2,2,2),
                           ellipse.level=0.99 #Confidence limits at 99%
                           )+
  scale_fill_manual(values=c("white","white","white"))+
  scale_colour_manual(values=c("red","black","blue"))+
  theme_bw()+
  ylim(c(-2.6,2.6))+
  xlab("Dimension 1: 54.9%")+
  ylab("Dimension 2: 25.7%")+
  theme_bw()+
  theme(legend.position = "bottom",
        plot.title = element_blank())

fviz.plot
```

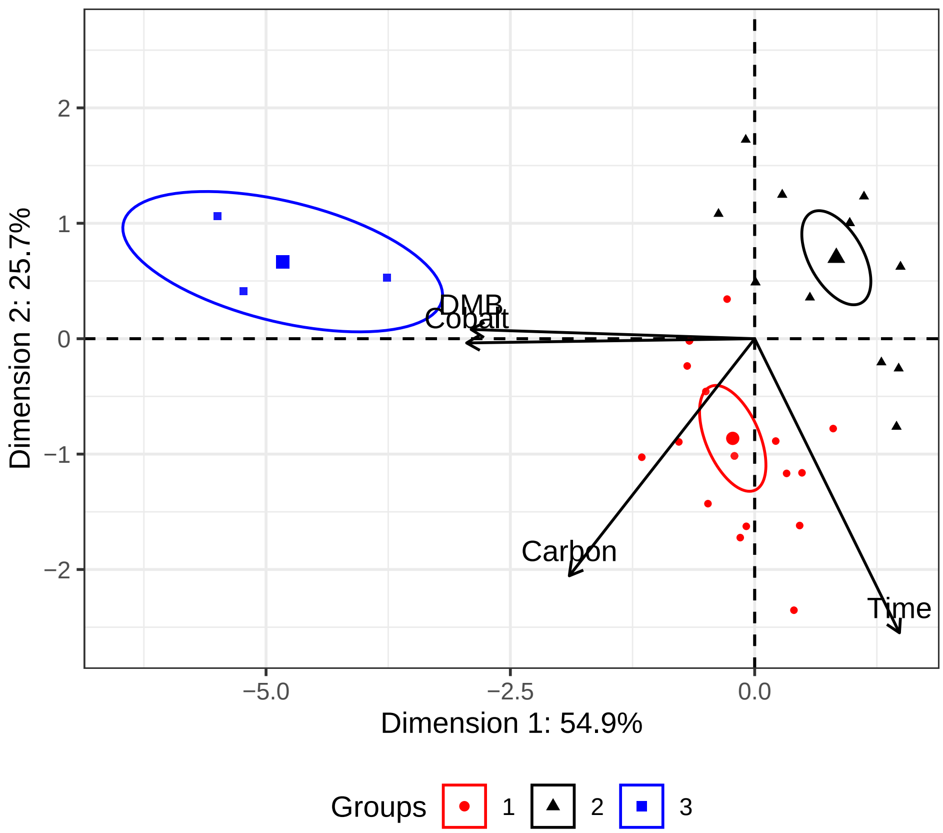


The PCA model generated was then used to intrapolate the relative position of the experimental conditions described in our experiments.

```
new_data <- data.frame(df[48,1:4])
new_data_centered <- scale(new_data, center = res.pca$center)
new_data_scores <- t(res.pca$rotation) %*% t(new_data_centered)
```

Lastly, the PCA including the intrapolated experimental point is visualised in the figure below.

```
fviz.plot2<-fviz.plot +
  geom_point(data = as.data.frame(t(new_data_scores)),
             aes(x = PC1, y = PC2),
             color = "purple", size = 2,
             shape=7, show.legend = FALSE)+ #Adding the point representing experimental conditions.
  geom_text(aes(label="E",x=-1.15,
                y=1.8, size=3), #Labeling the point as "E" for "Experimental"
            color="purple", show.legend = FALSE)

fviz.plot2
```

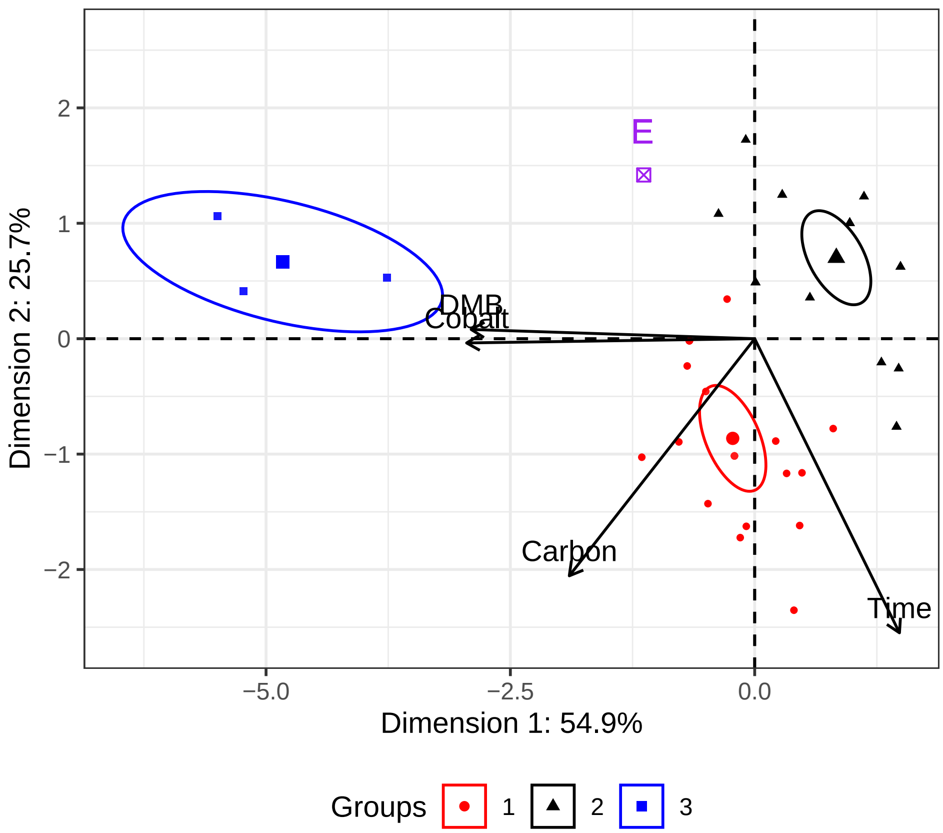

E
DMB
Cobalt
Carbon
Time
2
1
0
−1
−2
−5.0
−2.5
0.0
Dimension 1: 54.9%
Dimension 2: 25.7%
Groups
1
2
3

This is the information of the computer system in which the analysis was last performed to check output:

```r
sessionInfo()
```

```
## R version 4.2.3 (2023-03-15)
## Platform: x86_64-pc-linux-gnu (64-bit)
## Running under: Linux Mint 21.2
##
## Matrix products: default
## BLAS/LAPACK: /usr/lib/x86_64-linux-gnu/openblas-pthread/libopenblasp-r0.3.20.so
##
## locale:
##  [1] LC_CTYPE=en_IE.UTF-8       LC_NUMERIC=C
##  [3] LC_TIME=en_IE.UTF-8        LC_COLLATE=en_IE.UTF-8
##  [5] LC_MONETARY=en_IE.UTF-8    LC_MESSAGES=en_IE.UTF-8
##  [7] LC_PAPER=en_IE.UTF-8       LC_NAME=C
##  [9] LC_ADDRESS=C               LC_TELEPHONE=C
## [11] LC_MEASUREMENT=en_IE.UTF-8 LC_IDENTIFICATION=C
##
## attached base packages:
## [1] stats     graphics  grDevices utils     datasets  methods   base
##
## other attached packages:
##  [1] formatR_1.14     cluster_2.1.4    reshape2_1.4.4   factoextra_1.0.7
##  [5] lubridate_1.9.2  forcats_1.0.0    stringr_1.5.0    dplyr_1.1.1
##  [9] purrr_1.0.1      readr_2.1.4      tidyr_1.3.0      tibble_3.2.1
## [13] ggplot2_3.4.2    tidyverse_2.0.0
##
## loaded via a namespace (and not attached):
##  [1] tidyselect_1.2.0 xfun_0.38        carData_3.0-5    colorspace_2.1-0
##  [5] vctrs_0.6.1      generics_0.1.3   htmltools_0.5.5  yaml_2.3.7
##  [9] utf8_1.2.3       rlang_1.1.0      pillar_1.9.0     ggpubr_0.6.0
## [13] glue_1.6.2       withr_2.5.0      lifecycle_1.0.3  plyr_1.8.8
## [17] munsell_0.5.0    ggsignif_0.6.4   gtable_0.3.3     evaluate_0.20
## [21] labeling_0.4.2   knitr_1.42       tzdb_0.3.0       fastmap_1.1.1
## [25] fansi_1.0.4      highr_0.10       broom_1.0.4      Rcpp_1.0.10
## [29] scales_1.2.1     backports_1.4.1  abind_1.4-5      farver_2.1.1
## [33] hms_1.1.3        digest_0.6.31    stringi_1.7.12   rstatix_0.7.2
## [37] ggrepel_0.9.3    grid_4.2.3       cli_3.6.1        tools_4.2.3
## [41] magrittr_2.0.3   car_3.1-2        pkgconfig_2.0.3  timechange_0.2.0
## [45] rmarkdown_2.21   rstudioapi_0.14  R6_2.5.1         compiler_4.2.3
```

Please get in touch with the authors if you are having issues reproducing the code! All the best.
